# Discovery of compounds that inhibit SARS-CoV-2 Mac1-ADP-ribose binding by high-throughput screening

**DOI:** 10.1101/2022.03.01.482536

**Authors:** Anu Roy, Yousef M. Alhammad, Peter McDonald, David K. Johnson, Junlin Zhuo, Sarah Wazir, Dana Ferraris, Lari Lehtiö, Anthony K.L. Leung, Anthony R. Fehr

## Abstract

The emergence of several zoonotic viruses in the last twenty years, especially the pandemic outbreak of SARS-CoV-2, has exposed a dearth of antiviral drug therapies for viruses with pandemic potential. Developing a diverse drug portfolio will be critical for our ability to rapidly respond to novel coronaviruses (CoVs) and other viruses with pandemic potential. Here we focus on the SARS-CoV-2 conserved macrodomain (Mac1), a small domain of non-structural protein 3 (nsp3). Mac1 is an ADP-ribosylhydrolase that cleaves mono-ADP-ribose (MAR) from target proteins, protects the virus from the anti-viral effects of host ADP-ribosyltransferases, and is critical for the replication and pathogenesis of CoVs. In this study, a luminescent-based high-throughput assay was used to screen ∼38,000 small molecules for those that could inhibit Mac1-ADP-ribose binding. We identified 5 compounds amongst 3 chemotypes that inhibit SARS-CoV-2 Mac1-ADP-ribose binding in multiple assays with IC_50_ values less than 100*µ*M, inhibit ADP-ribosylhydrolase activity, and have evidence of direct Mac1 binding. These chemotypes are strong candidates for further derivatization into highly effective Mac1 inhibitors.

## INTRODUCTION

COVID-19, caused by severe acute respiratory syndrome coronavirus 2 (SARS-CoV-2), is one of the most disruptive and deadly pandemics in modern times, with greater than 385 million cases and having led to greater than 5.7 million deaths worldwide. SARS-CoV-2 is the third CoV to emerge into the human population in the last 3 decades, following outbreaks of SARS-CoV in 2002-2003 and Middle East respiratory syndrome MERS-CoV in 2012. These outbreaks highlight the potential for CoVs to cross-species barriers and cause severe disease in a new host. There is a tremendous need to develop broad-spectrum antiviral therapies capable of targeting a wide range of CoVs to prevent severe disease following zoonotic outbreaks.

Coronaviruses encode for 16 highly conserved, non-structural proteins that are processed from two polyproteins, 1a and 1ab (pp1a and pp1ab) (1). The largest non-structural protein is non-structural protein 3 (nsp3) that encodes for multiple modular protein domains. Both the SARS-CoV and the SARS-CoV-2 nsp3 proteins include three tandem macrodomains, Mac1, Mac2, and Mac3 (2). Mac1 is present in all CoVs, unlike Mac2 and Mac3, and contains a conserved three-layered α/β/α fold, a common feature amongst all macrodomains. All CoV Mac1 proteins tested have mono-ADP-ribosylhydrolase (ARH) activity, though it remains unclear if they have significant poly-ARH activity (3-8). In contrast, Mac2 and Mac3 fail to bind ADP-ribose and instead bind to nucleic acids (9,10). Mac1 homologs are also found in alphaviruses, Hepatitis E virus, and Rubella virus, indicating that ADP-ribosylation may be a potent anti-viral post-translational modification (PTM) (11,12). All are members of the larger MacroD-type macrodomain family, which includes human macrodomains Mdo1 and Mdo2 (13).

ADP-ribosylation is a post-translational modification catalyzed by ADP-ribosyltransferases (ARTs, also known as PARPs) through transferring an ADP-ribose moiety from NAD^+^ onto target proteins or nucleic acids (14). ADP-ribose is transferred in as a single unit as mono-ADP-ribose (MAR), or it is transferred consecutively and covalently attached through glycosidic bonds to preceding ADP-ribose units to form a poly-ADP-ribose (PAR) chain. Both mono- and poly-ARTs inhibit virus replication, implicating ADP-ribosylation in the host-response to infection (15).

Several reports have addressed the role of Mac1 on the replication and pathogenesis of CoVs, mostly using the mutation of a highly conserved asparagine to alanine (N41A-SARS-CoV). This mutation abolished the MAR-hydrolase activity of SARS-CoV Mac1 (16). This mutation has minimal effects on CoV replication in transformed cells, but reduces viral load, leads to enhanced IFN production, and strongly attenuates both murine hepatitis virus (MHV) and SARS-CoV in mouse models of infection (4,16-18). Murine hepatitis virus strain JHM (MHV-JHM) Mac1 was also required for efficient replication in primary macrophages, which could be partially rescued by the PARP inhibitors or siRNA knockdown of PARP12 or PARP14 (19). These data suggest that Mac1’s function is to counter PARP-mediated anti-viral ADP-ribosylation (20). More recently, we have identified mutations in the MHV-JHM Mac1 domain, predicted to abolish ADP-ribose binding, that resulted in severe replication defects in cell culture, indicating that for some CoVs Mac1 may be even more important than previously appreciated (21). Mutations in the alphavirus and HEV macrodomain also have substantial phenotypic effects on virus replication and pathogenesis (22-26).

As viral macrodomains are critical virulence factors, they are unique targets for anti-viral therapeutics (20). Several studies have reported structures that could potentially bind to the ADP-ribose binding pocket of SARS-CoV-2 Mac1. While most of these studies were limited to *in silico* studies, a few have tested compound activity in biochemical assays, but have been met with minimal success (27-30). The only compounds identified thus far that inhibit SARS-CoV-2 Mac1 with IC_50_ less than 100 μM are Suramin, which inhibited Mac1-ADP-ribose binding in a FRET assay with an IC_50_ of 8.7 μM, and Dasatanib, which inhibited Mac1 mono-ARH activity with an IC_50_ of ∼50 μM. Suramin targeted several divergent macrodomains and is known to have additional targets, and thus is not suitable for further evaluation (30). Dasatinib is not a candidate for a Mac1 inhibitor as it is toxic to mammalian cells, though it may provide a scaffold for further inhibitor development. None of the identified compounds have been tested for their ability to inhibit Mac1 in cell culture or in animal models of disease.

Here, we optimized two high-throughput macrodomain-ADP-ribose binding assays, a previously described luminescent-based AlphaScreen™ assay, and a novel fluorescence polarization assay (31,32), and used the AlphaScreen™ assay to screen ∼38,000 compounds for their ability to inhibit SARS-CoV-2 Mac1-ADP-ribose binding. We identified 5 compounds from 3 chemotypes that inhibited ADP-ribose binding by the SARS-CoV-2 Mac1 protein in both assays, some with IC_50_ values as low as 5-10 μM. These compounds also demonstrated some inhibition of ARH activity and have evidence of direct binding to Mac1. The profiling of the most potent inhibitor against a panel of virus and human MAR binding and hydrolyzing proteins revealed the remarkable selectivity of the inhibition of SARS-CoV-2 Mac1. These compounds represent several series that can be further developed into potent Mac1 inhibitors and potential therapeutics for SARS-CoV-2 and other CoVs of interest.

## RESULTS and DISCUSSION

### Comparison of viral and human macrodomains in two high-throughput ADP-ribose binding assays

Here we established two distinct ADP-ribose binding assays for multiple macrodomain proteins (Fig. 1A-C). First, we adopted a previously published AlphaScreen™ (AS) assay, where a short peptide was modified at a leucine residue with ADP-ribose through an amino-oxyacetic acid linkage, and at a second leucine residue with biotin (Fig. 1A) (32). Streptavidin donor beads and Ni^2+^ acceptor beads induce a light signal if the His-tagged Mac1 protein interacts with the biotinylated peptide (Fig. 1B). We also developed a fluorescent polarization (FP) assay as an orthogonal assay to evaluate interactions of macrodomains with ADP-ribosylated peptide. This assay used the same peptide but with fluorescein attached instead of biotin and measures polarization of the fluorescent signal (Fig. 1C). We then tested 4 separate macrodomains for their ability to bind to these peptides, the human macrodomain Mdo2, and Mac1 from SARS-CoV, MERS-CoV, and SARS-CoV-2. All 4 macrodomains bound to the ADP-ribosylated control peptides better than to non-ADP-ribosylated peptides (Fig. 1D,G). The AS assay had an especially strong signal-to-background ratio, ranging from ∼0.75-2×10^3^. To further study the binding of Mac1 proteins to AS and FP peptides, we evaluated binding in a dose-dependent assay. Of these four proteins, the human MDO2 demonstrated the highest affinity in both assays, with a K_D_ of 1.1 ± 0.3 µM in the FP assay and reached a maximum signal in the AS assay at 40 nM (Fig. 1E,H). The SARS-CoV-2 Mac1 had a K_D_ of 3.4 ± 0.4 µM in the FP assay and reached a maximum signal in the AS assay at 0.625 µM, while the SARS-CoV and MERS-CoV Mac1 both reached their maximum signal in the AS assay at ∼1.25-2.5 µM (AS) and had K_D_’s of 7.7 ± 1.3 µM and 19.9 ± 3.3 µM in the FP assay, respectively (Fig. 1F,I).

**Figure 1.**
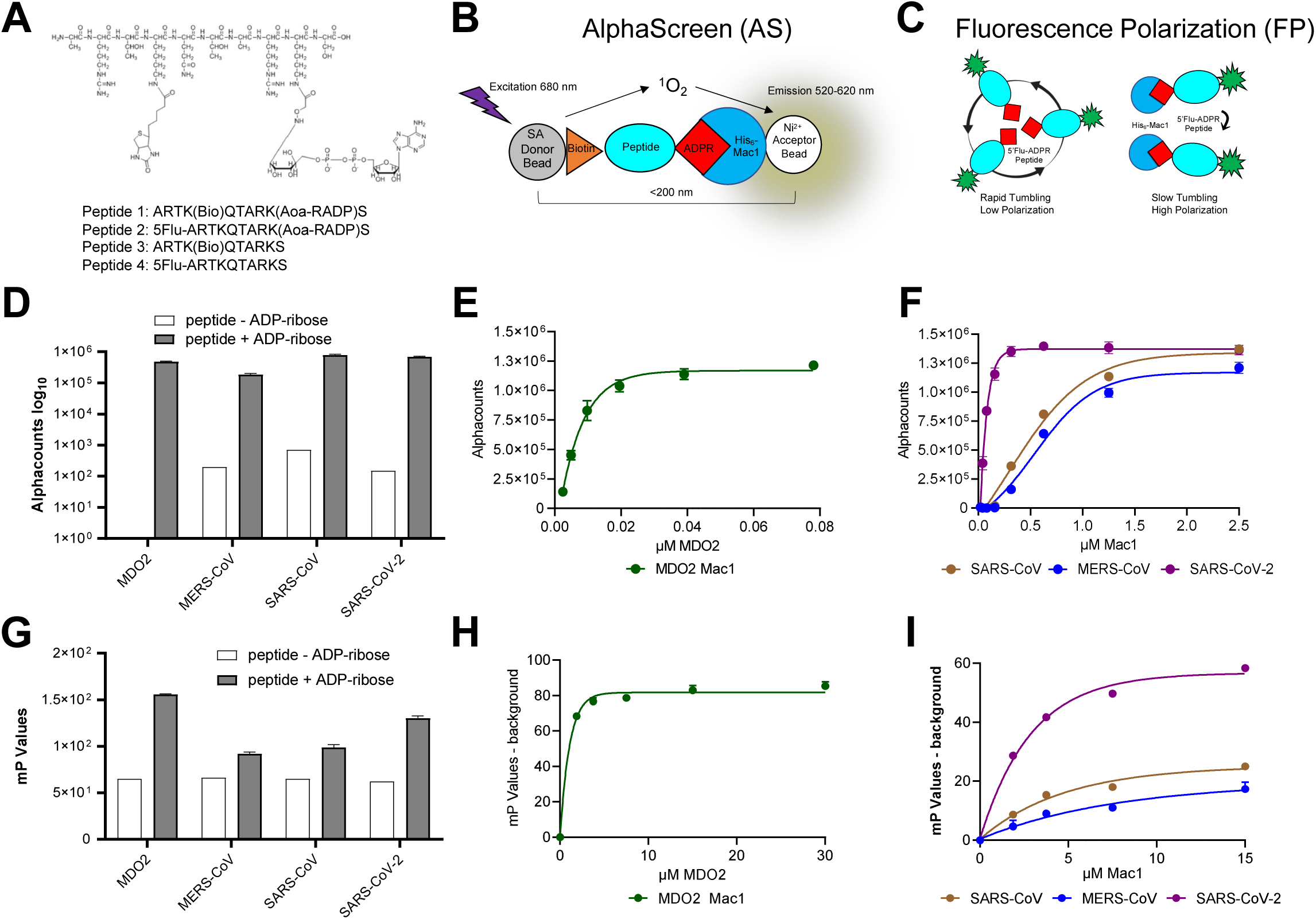
Coronavirus Mac1 binding to ADP-ribosylated peptides. A) Illustration of the amino-oxyacetic acid modified lysine-conjugated ADP-ribosylated peptide with an additional biotin conjugated to a different lysine residue and included are the amino acid sequences and modification sites of peptides used in this study. B-C) Cartoon diagrams depicting a bead-based AS (A) and FP (B) assays for measuring macrodomain interactions with an ADP-ribosylated peptide. D) Macrodomain proteins were incubated with peptide #1 or peptide #3 for 1 hour at RT and Alphacounts were determined as described in Methods. E-F) Peptide #1 was incubated with indicated macrodomains at increasing concentrations and Alphacounts were measured as previously described. G) Mac1 proteins were incubated at indicated concentrations with peptide #2 or peptide #4 and the plate was incubated at 25°C for 1 hr before polarization was determined. H-I) Peptide #2 was incubated with indicated macrodomain proteins at increasing concentrations and polarization was determined as previously described. All data represent the means ± SD of 2 independent experiments for each protein.

Next, we tested the ability of free ADP-ribose to inhibit the binding of Mac1 to the ADP-ribosylated peptide. For these displacement assays, the amount of beads, peptide, and Mac1 protein amounts to be used were optimized to obtain a robust signal while limiting the amount of reagents used for screening purposes (see Methods). The addition of free ADP-ribose, but not ATP, into the AS and FP assays inhibited human macrodomain and CoV Mac1 binding to the ADP-ribosylated peptides, confirming that these assays can be used to identify macrodomain binding inhibitors (Fig. 2). IC_50_ values for free ADP-ribose ranged between 0.24 µM with SARS-CoV Mac1 to 1.5 µM with SARS-CoV-2 using the free ADP-ribose in the AS assay (Fig. 2A). Similar results, albeit higher IC_50_ values were observed in the FP assay, likely because of higher amount of Mac1 used in this assay (4 µM vs 250 nM), with IC_50_ values ranging from 2.3 µM to 9.74 µM (Fig. 2B).

**Figure 2.**
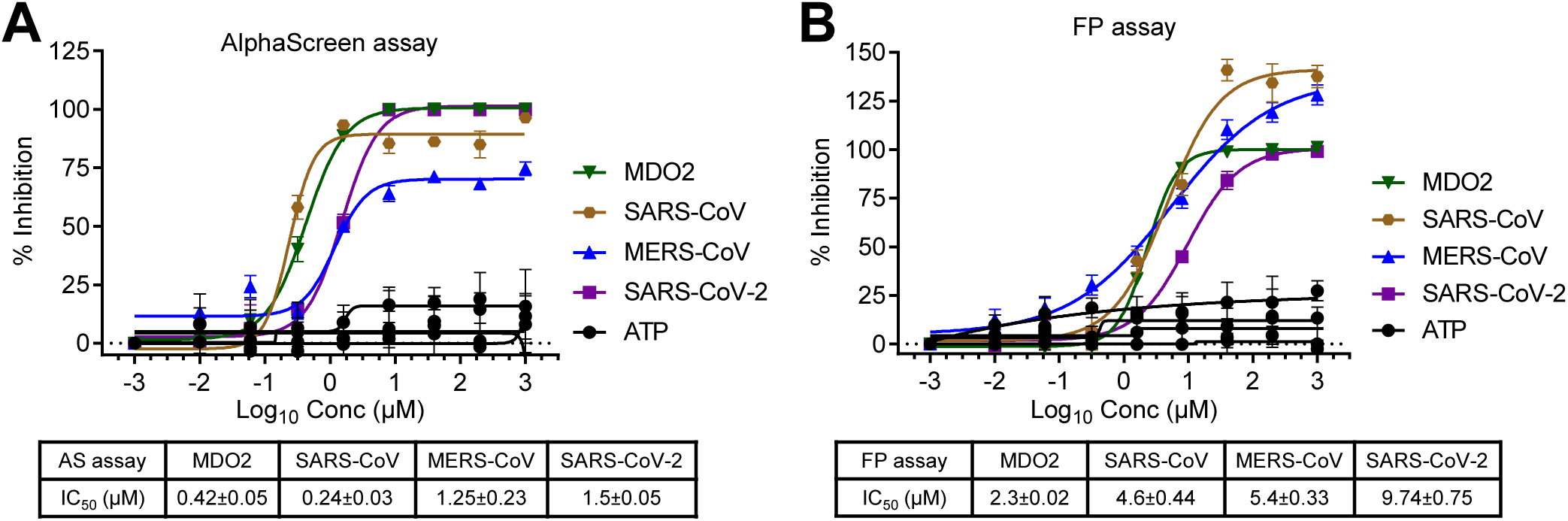
Free ADP-ribose inhibits macrodomain binding to ADP-ribosylated peptides. ADP-ribose competition assays were used to block the interaction between macrodomain proteins and ADP-ribosylation peptides in the AS (A) or FP (B) assays. ATP was used as a negative control. The data represent the means ± SD of 2 independent experiments for each protein.

### High-throughput screening (HTS) for SARS-CoV-2 Mac1 inhibitors

We next performed a small pilot screen of ∼ 2,000 compounds from the Maybridge Mini Library of drug-like scaffolds at 10 µM using both AS and FP assays (Fig. 3A-B). We identified 39 compounds that significantly inhibited Mac1-ADP-ribose binding at >3 standard deviations (3SD) plus the plate median (Fig. 3A-B). After performing dose-response curves we found that two compounds inhibited binding in both assays (Fig. 4A). We then tested these compounds in a counter screen, which is also an AS assay that utilizes a biotinylated-His peptide that gives off a strong signal with the addition of streptavidin donor and nickel acceptor beads. These two compounds did not affect the signal from our counter screen indicating that they do not intrinsically inhibit the assay. After this initial validation of our screen, three additional libraries were chosen to include a total number of 35,863 compounds from the Analyticon, 3D BioDiversity, and Peptidomimetics libraries (Fig. 3A). We chose the AS assay as our primary HTS assay, as the average Z’ score for the AS was higher than the Z’ score from the FP assay in our original screen (0.82 vs 0.67). In this larger screen, the average Z’ was 0.89±0.05, indicating a strong separation between positive and negative controls (Fig. 3C). Using the same hit criteria described above for each individual library, we identified 406 hits resulting in a 1% hit rate (Fig. 3D). Of note, the Analyticon library produced a lot of non-specific inhibitors, indicating a lot of these compounds likely inhibit the assays themselves (Fig. 3B). We next performed dose-response (10-40 µM) curves of these 406 compounds in our primary (AS), orthogonal (FP), and counter screen (Bn-His_6_) assays (Fig. 3). From the 406 original hits, 26 compounds were identified that inhibited SARS-CoV-2 Mac1-ADP-ribose binding in the AS assay in a dose-dependent fashion, and 6 compounds were identified that inhibited Mac1 binding in both AS and FP assays (Fig. 3D). Of these 32 hit compounds, we re-purchased 17 of them, excluding 15 based on several selection criteria, including substantial inhibition of the counter screen, high IC_50_ values in the AlphaScreen, pan-assay interference compounds, and compound availability (Fig. 3D). The remaining 17 compounds along with 4 analogs were repurchased or resynthesized (see Methods).

**Figure 3.**
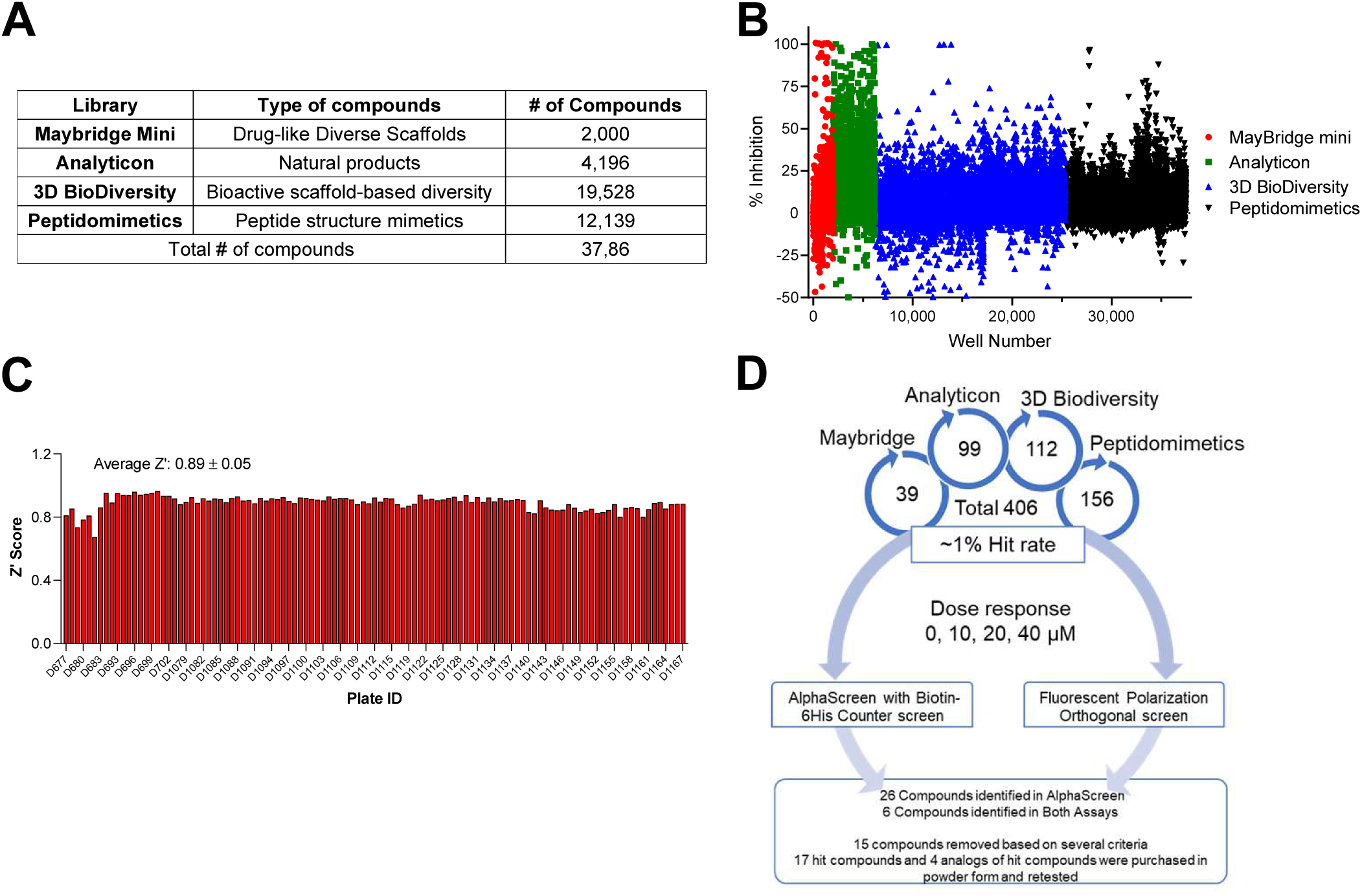
High-throughput screen for SARS-CoV-2 Mac1 inhibitors. A) List of libraries that were screened, the number of compounds from each library, and the type of compounds each library contains. B) Scatterplot showing the % inhibition of each compound in the screen. The cutoff for a hit was the plate median + 3 standard deviations. C) Z’ scores were determined for each plate in the screen. The average Z’ score was 0.89 ± 0.05. D) Dose response confirmation. From the original screen, we identified 406 potential hits, these hits were retested in a dose-response assay on both the AS and FP assays and were also counterscreened against a biotinylated 6His peptide. After these assays and other exclusion criteria, 17 hit compounds and 4 analogs were repurchased or resynthesized.

**Figure 4.**
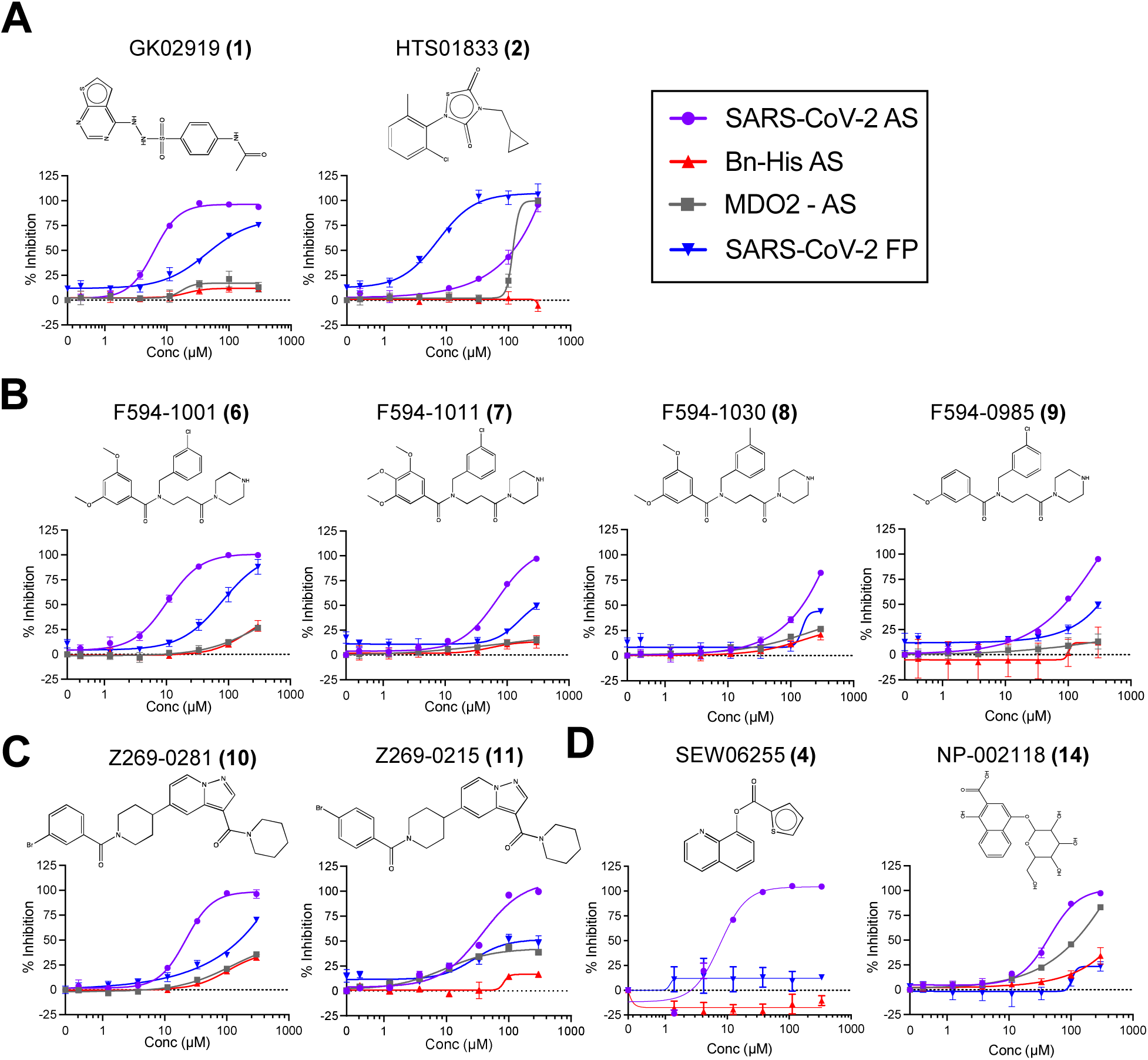
Identification of chemical compounds that inhibit SARS-CoV-2 Mac1 ADP-ribose binding. Dose-response curves representing hit compounds identified in the HTS. A) Maybridge Mini Library compounds **1, 2**. B) Compound **6** and its analogs, **7, 8, 9**. C) Compound **10** and its analog **11**. D) Compounds **4** and **14** which did not inhibit FP assay. Data represent the means ± SD of at least 2 independent experiments for each protein. Structures were created using ChemDraw.

Re-purchased compounds were evaluated in dose-response assays against both SARS-CoV-2 Mac1 and human MDO2 protein. Our cutoff criteria included: *i)* compound must inhibit both primary and orthogonal assays with at least 75% inhibition in AS assay and at or near 50% inhibition in the FP assay, and *ii)* less than 30% inhibition of the Bn-His_6_ counter screen. Among the 17 selected and the 4 analogs compounds, six compounds inhibited ADP-ribose binding of SARS-CoV-2 Mac1 in both AS and FP assays with no substantial inhibition of the Bn-His_6_ counter screen. These were compounds **1**,**2**,**6**,**7**,**10**, and **11** (Table 1). IC_50_ values ranged from 6.2 µM to 112.2 µM in AS assay and 7.3 µM to 159.4 µM in FP assay (Table 1, Fig. 4). Compounds **1, 10**, and **11** also had some inhibitory activity against the MERS-CoV Mac1 protein, though the inhibition of MERS-CoV Mac1 was lower than the inhibition demonstrated against SARS-CoV-2 (Table 1). In addition, only compound **2** inhibited MDO2, indicating that these compounds were broadly specific for viral macrodomains.

**Table 1:**
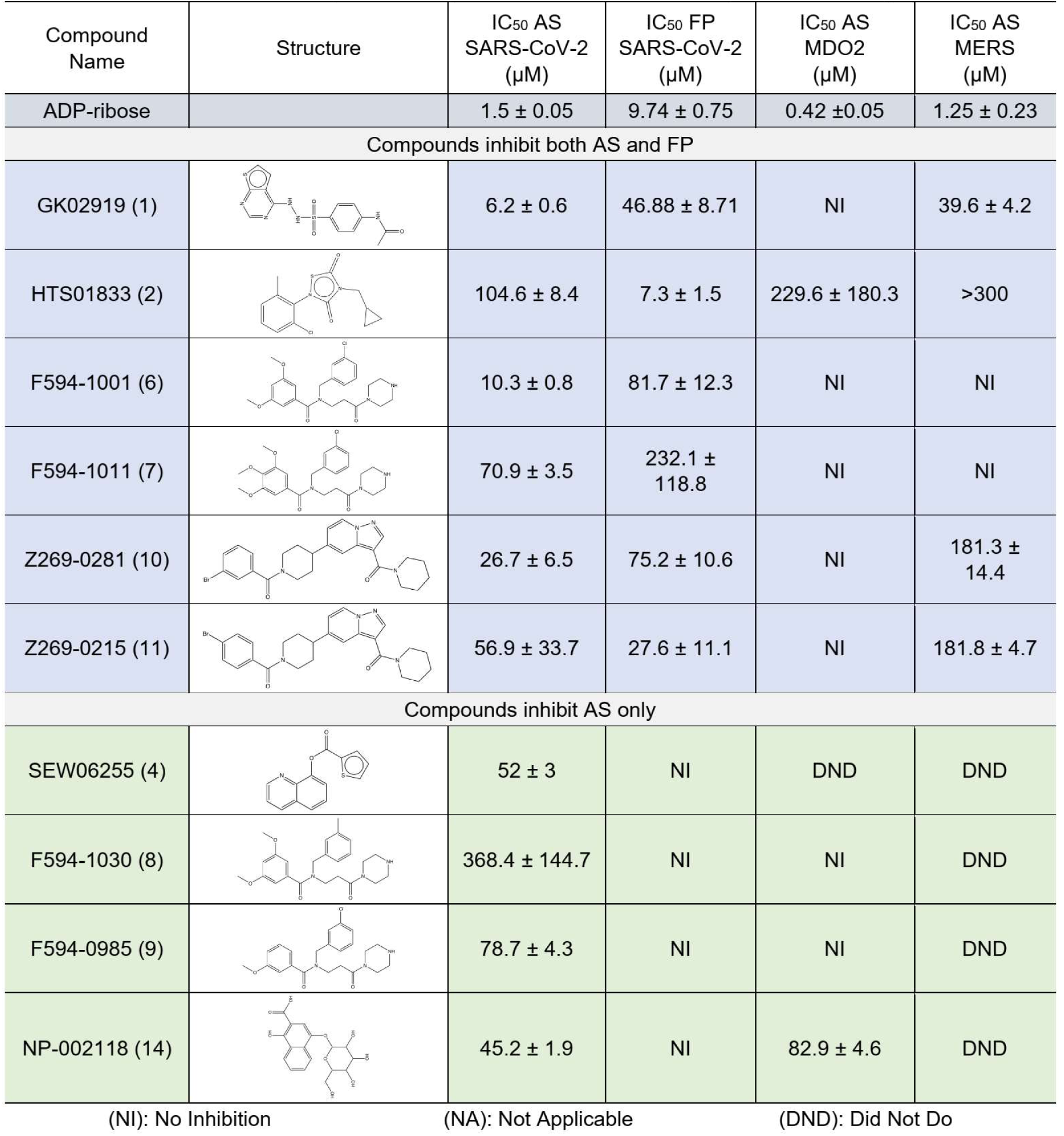
IC_50s_ of the selected compounds in AlphaScreen and Fluorescence polarization assays

| Compound Name | Structure | IC<sub>50</sub> AS SARS-CoV-2 (μM) | IC<sub>50</sub> FP SARS-CoV-2 (μM) | IC<sub>50</sub> AS MDO2 (μM) | IC<sub>50</sub> AS MERS (μM) |
| --- | --- | --- | --- | --- | --- |
| ADP-ribose |  | 1.5 ± 0.05 | 9.74 ± 0.75 | 0.42 ± 0.05 | 1.25 ± 0.23 |
| Compounds inhibit both AS and FP |  |  |  |  |  |
| GK02919 (1) |  | 6.2 ± 0.6 | 46.88 ± 8.71 | NI | 39.6 ± 4.2 |
| HTS01833 (2) |  | 104.6 ± 8.4 | 7.3 ± 1.5 | 229.6 ± 180.3 | >300 |
| F594-1001 (6) |  | 10.3 ± 0.8 | 81.7 ± 12.3 | NI | NI |
| F594-1011 (7) |  | 70.9 ± 3.5 | 232.1 ± 118.8 | NI | NI |
| Z269-0281 (10) |  | 26.7 ± 6.5 | 75.2 ± 10.6 | NI | 181.3 ± 14.4 |
| Z269-0215 (11) |  | 56.9 ± 33.7 | 27.6 ± 11.1 | NI | 181.8 ± 4.7 |
| Compounds inhibit AS only |  |  |  |  |  |
| SEW06255 (4) |  | 52 ± 3 | NI | DND | DND |
| F594-1030 (8) |  | 368.4 ± 144.7 | NI | NI | DND |
| F594-0985 (9) |  | 78.7 ± 4.3 | NI | NI | DND |
| NP-002118 (14) |  | 45.2 ± 1.9 | NI | 82.9 ± 4.6 | DND |

(NI): No Inhibition
(NA): Not Applicable
(DND): Did Not Do

**Table 2:** Peak values of DSF thermal shift temperatures

| | Peak $\Delta T_m$ | | | | | |
| --- | --- | --- | --- | --- | --- | --- |
| **Compound** | **1** | **2** | **6** | **7** | **10** | **11** |
| **Average** | 0.68 | 18.88- | 1.67 | 0.22 | 0.51 | 0.47 |
| **SD** | 0.04 | 0.53 | 0.21 | 0.08 | 0.03 | 0.04 |
| **Conc ($\mu\text{M}$)** | 150 | 300 | 300 | 300 | 150 | 150 |

### Selected compounds demonstrate evidence of SARS-CoV-2 Mac 1 binding

Next, we set out to test the hypothesis that these compounds inhibit Mac1-ADP-ribose by binding to Mac1, and not other components of the assay, such as the peptide. To test for Mac1 binding, we used a differential scanning fluorimetry (DSF) assay as previously described (8) and tested our top 6 hit compounds (Fig. 5, S1) and compounds **8** and **9**, as they are analogs of **6** and **7** (Fig. S2). In this assay, compound binding to Mac1 should increase the melting temperature of Mac1. The addition of free ADP-ribose, which binds to Mac1, showed a dose-dependent increase of approximately 4஬ in the melting temperature of Mac1, while the negative control, ATP, had no effect, as previously demonstrated (8). **1, 6, 7, 10**, and **11** showed dose-dependent shifts in the melting temperature of Mac1 ranging from 0.2 - 1.5஬, providing strong evidence that these compounds bind to Mac1, albeit not with the same affinity as ADP-ribose. On the other hand, compound **2** resulted in highly irregular thermal shift curves, indicating that this compound may not be a true Mac1 binder (Fig. 5, S1). These results provide evidence that 5 of our 6 hit compounds (**1, 6, 7, 10**, and **11**) directly bind to SARS-CoV-2 Mac1.

**Figure 5.**
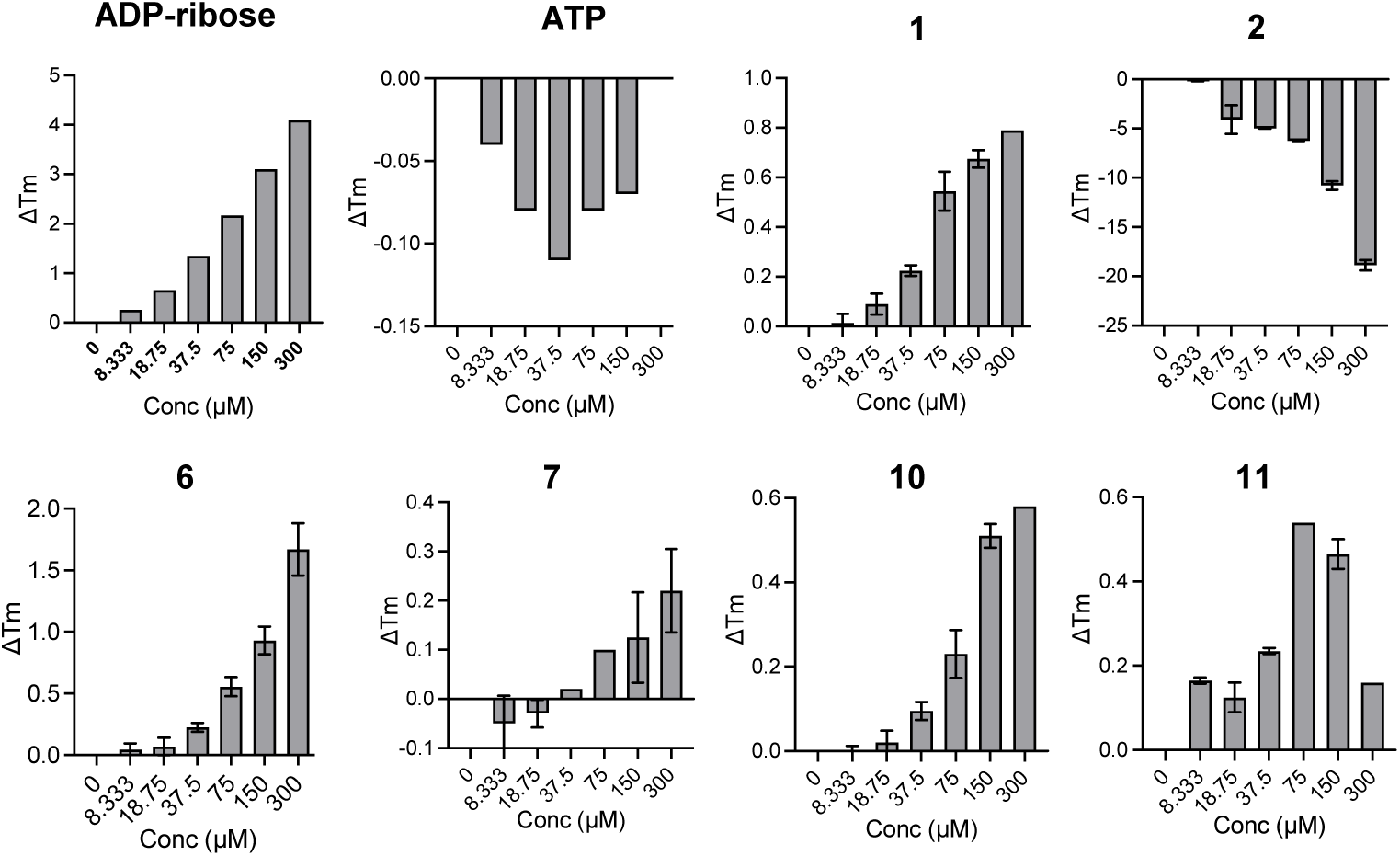
Thermal stability of SARS-CoV-2 Mac1 after incubation with hit compounds. The top 6 hit compounds were tested for their ability to increase the thermal stability of SARS-CoV-2 Mac1 in a differential scanning fluorimetry assay (DSF). The data represent the means ± SD of the ΔT_m_ from two independent experiments.

### Hit compounds inhibit ADP-ribosylhydrolase activity *in vitro*

SARS-CoV-2 Mac1 is a mono-ADP-ribosylhydrolase that removes mono-ADP-ribose from target proteins (8). Next, we examined the ability of some of our top 5 hit compounds to inhibit the enzymatic activity of SARS-CoV-2 Mac1 using two distinct assays. The first approach was a gel-based Mac1 ADP-ribosylhydrolase assay where we tested each compound against the SARS-CoV-2 Mac1 protein (8). Compound **1** tended to precipitate in these assays at higher concentrations, and so we used lower concentrations for this compound than others. Compounds **1, 6**, and **7** exhibited a dose-dependent inhibition of Mac1 ADP-ribosylhydrolase activity (Fig. 6A). We were unable to detect any significant inhibition with **10** and **11** in this assay.

**Figure 6.**
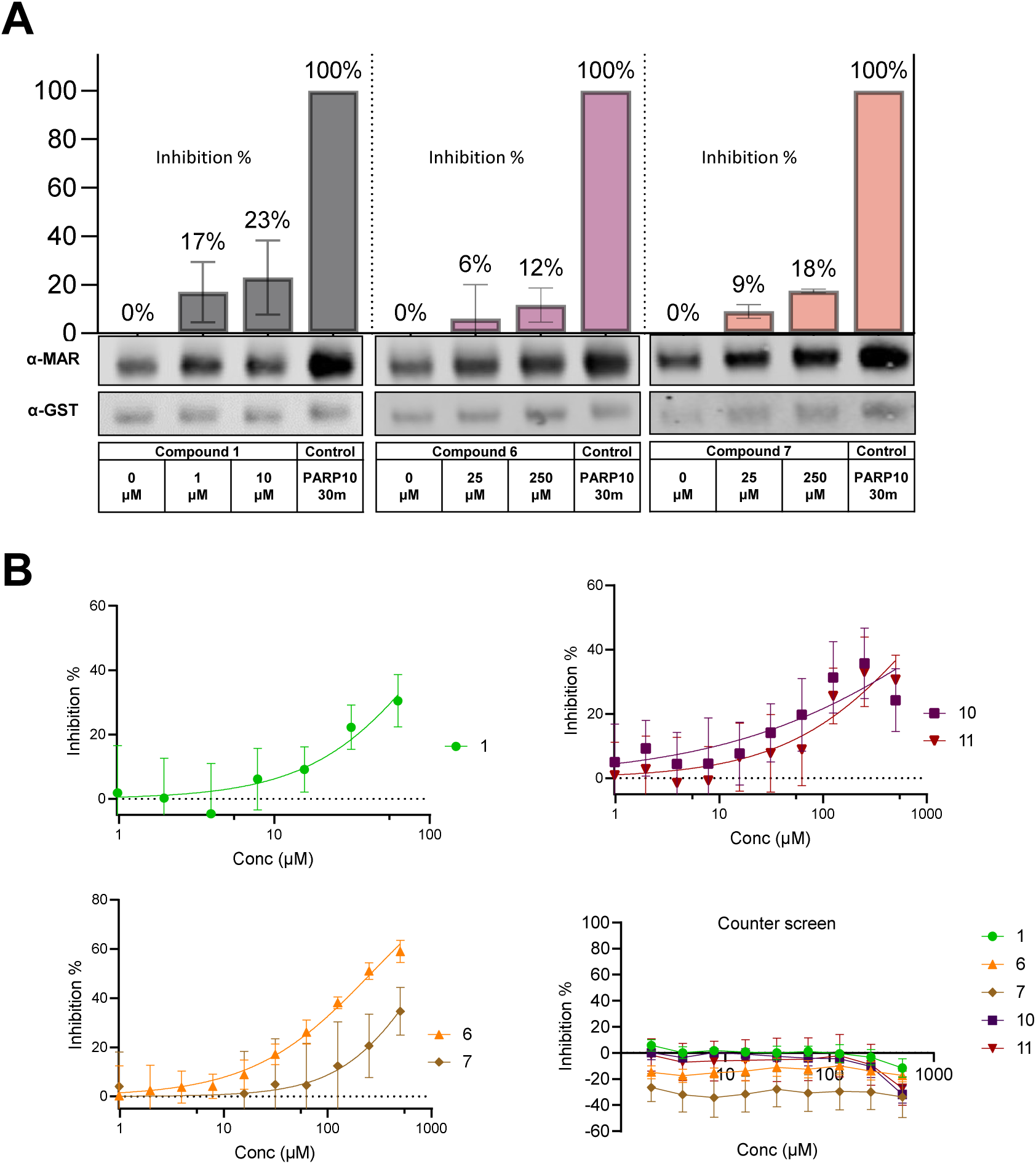
Impact of hit compounds on SARS-CoV-2 ADP-ribosylhydrolase activity. A) Compounds were incubated at indicated concentrations for 30 minutes with the SARS-CoV-2 Mac1 protein prior to adding the PARP10 substrate and then were further incubated for 30 minutes. Proteins were analyzed by Immunoblotting with anti-GST (PARP10) and anti-MAR binding reagent (MABE1076). Gels were quantitated using Image Studio software. The bar graph above each immunoblots represent the mean inhibition ± SD from at least two independent experiments.

Next, we utilized a recently published high-throughput luminescence-based ADP-ribosylhydrolyase assay (33). Here we found that **1, 6, 7, 10** and **11** all showed dose-dependent inhibition of ADP-ribosylhydrolase activity (Fig. 6B). **6** was clearly the most efficient inhibitor, as it had a peak of ∼60% inhibition, similar to Dasatinib which we previously identified in a separate HTS (33). In contrast to the gel-based assay, **10** and **11** did inhibit ADP-ribosylhydrolase activity in this assay, likely reflecting the increased sensitivity of this assay compared to the gel-based assay. These results indicate that the identified Mac1 inhibitors block Mac1 binding and Mac1 enzymatic activity.

### Selectivity Profiling

As compound **6** inhibited both Mac1 ADP-ribose binding and hydrolysis activity, and showed the strongest evidence of direct Mac1 binding, we tested its ability to inhibit 16 different macrodomains using a recently developed FRET-based assay (30). Again, **6** demonstrated dose-dependent inhibition of Mac1-ADP-ribose binding in this assay, consistent with our AS results but with a slightly higher IC_50_ of 45.0 ± 10.9 µM (Fig. 7A). Remarkably, when tested again 16 different human and viral macrodomains in this assay, **6** only inhibited SARS-CoV-2 Mac1, having only minimal levels in inhibition of all other macrodomain proteins, including other CoV macrodomains (Fig. 7B), which is in agreement with the selectivity observed in AS (Table 1). These results indicate that this compound is highly selective for the SARS-CoV-2 Mac1 protein.

**Figure 7.**
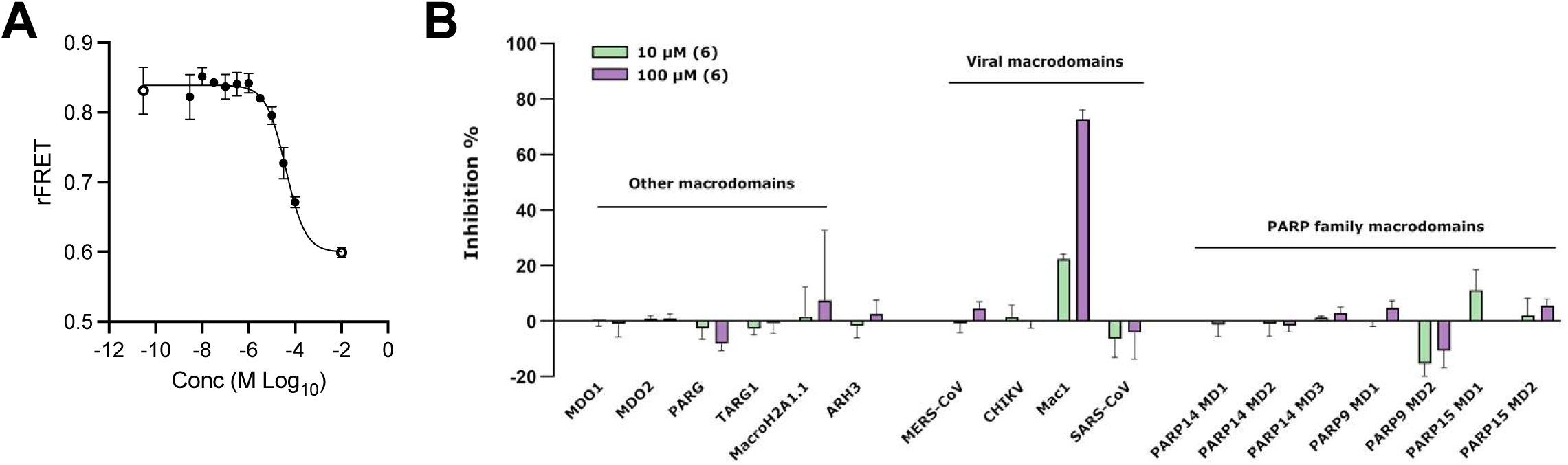
Compound 6 is highly selective for the SARS-CoV-2 Mac1 protein. A-B) Compound **6** was tested in a FRET-based assay for its ability to inhibit SARS-CoV-2 Mac1 protein in a dose-dependent manner (A) and for its ability to inhibit a panel of 17 macrodomain containing proteins (B). The data in means ± SD are shown as a single experiment representative of 3 independent experiments.

### Structure activity relationship (SAR)

The top 5 compounds could be separated into 3 chemotypes based on their structures. To analyze the involved residues and type of connection between selected hit compound and Mac1, we used computational docking analysis to get an initial structure activity relationship (SAR) by predicting poses of compounds in Mac1 structures. In addition to our 5 hit compounds, we also docked compounds **8** and **9** as they are analogs of **6** and **7** and could give further insight into SAR, even though we either detected minimal or no direct Mac1 binding by these compounds. These seven compounds were docked against the ADP-ribose bound structure of SARS-CoV-2 Mac1 (PDB 6WOJ) as well as three apo structures of Mac1 were used (PDB 7KR0, 7KR1, 6WEY). Docking and glide emodel scores were calculated for each compound against all four structures and the best structure was chosen based on these scores (Table S1). Analog compounds **6, 7, 8**, and **9** were assessed both based on score and visual inspection, and were re-docked using a core constraint to a high scoring, intuitive pose of compound **7**. All top scoring poses were subsequently minimized using Prime, allowing flexibility within 5 Å of the ligand. Compound **1** was its own chemotype but has a sulfonohydrazide that is also found in a compound identified in a previous screen for Mac1 compounds (34). It also has a thienopyrimidine that is similar to the pyrrolopyrimidine found in of the compounds identified in the fragment screen by Schuller et al (31). It makes a hydrogen bond with a backbone amine of D22, pie-stacking interactions with F156, and extends with a benzene ring into the distal ribose pocket inserting in between the GGG and GIF loops (Fig. 8A). Compounds **10** and **11** are close analogs with a single difference of positioning in the bromobenzoyl moiety on the piperidine ring (Fig. 4C). These compounds had similar activity across the board in our assays, making it difficult to analyze their SAR. While they docked into the binding pocket, these docking poses only indicate a single hydrogen bond with the backbone amino of D22. In contrast, compounds **6, 7, 8**, and **9** are close analogs of each other and have a wide-range of inhibitory and binding activity. IC_50_ values for these compounds range from 10 to several hundred µM (Table 1). Direct binding also varied substantially, with T_m_’s ranging from ∼1.7 ஬ (**6**) to undetectable binding (**8**). These compounds all have the same base structure, including a beta-alanine core substituted with a N-benzyl or N-chlorobenzyl group, a methoxy benzoyl group and a piperazine amide. The main difference between **6** and its analogs are the addition of a methoxy group on the benzoyl group (**7**), the loss of a chlorine (**8**), and a missing methoxy group (**9**). Each of these changes reduces the activity of this series indicating that *i)* the orientation of the methoxy groups on **6** is likely important for its increased activity, *ii*) reorienting **7** to accommodate the 4-methoxy group likely decreases activity due to the disruption of multiple interactions, and *iii)* the chlorine likely makes a critical halogen bond with a backbone amino group of L126 in the binding pocket.

**Figure 8.**
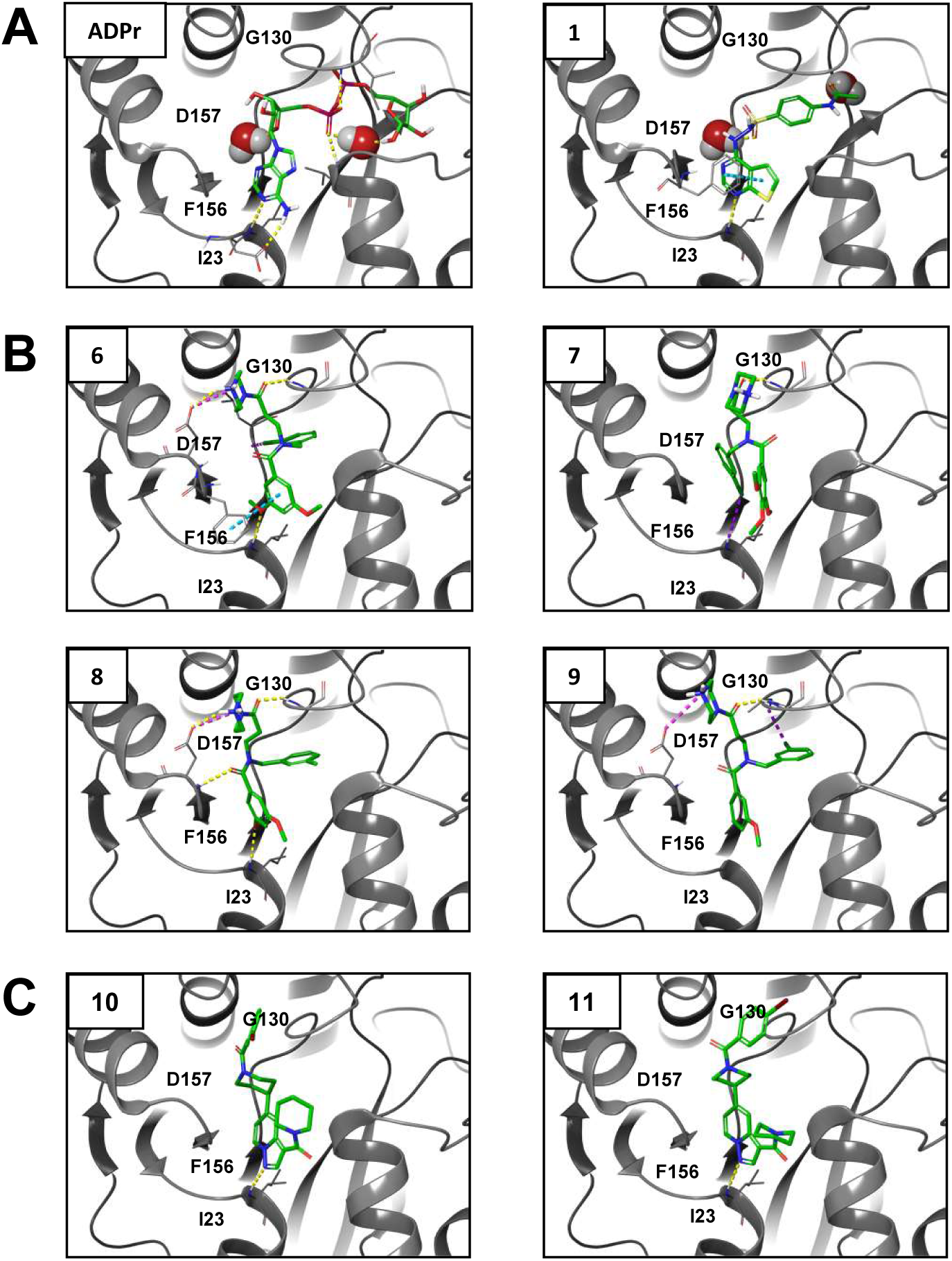
Computational modeling of identified compounds with SARS-CoV-2 Mac1 structures. Indicated compounds were docked and modeled with SARS-CoV-2 Mac1 structures using Maestro Schrödinger software and separated into 3 groups. (A) – Compound 1; (B) Compounds 6, 7, 8, 9; (C) Compounds 10, 11. Yellow lines - hydrogen bonds; Cyan lines - pi-pi interactions; magenta lines – weak hydrogen bonds; and purple lines – halogen bond.

In conclusion, we developed multiple high-throughput ADP-ribose binding assays and performed HTS to identify high-quality Mac1 inhibitors. We followed these screens with several additional assays to measure their ability to inhibit ADP-ribosylhydrolase activity and their direct binding to Mac1. We have identified 5 compounds that inhibit both the primary and orthogonal assays without inhibiting the counter screen and demonstrate dose-dependent inhibition of Mac1 enzymatic activity. Compounds **1** and **6** are particularly effective with IC_50_ values of ∼10 µM in the AS assay, along with thermal shifts and docking poses that indicate direct binding to Mac1. Compound **6** shows excellent selectivity towards SARS-CoV-2 over the human macrodomains guiding further development of the compound. We expect that these compounds could be utilized for further derivatization and optimization into more potent Mac1 inhibitors.

## METHODS

### Reagents

All plasmids and proteins used were expressed and purified as previously described (30,35-37). All compounds were repurchased from MolPort except for compounds **6** and **10**, which were repurchased from ChemDiv. After reordering once, compounds **10** and **11** became unavailable and thus were resynthesized according to the literature (38). ADP-ribosylated peptides were purchased from Cambridge peptides.

### Differential Scanning Fluorimetry (DSF)

Thermal shift assay with DSF involved use of LightCycler® 480 Instrument (Roche Diagnostics). In total, a 15 μL mixture containing 8X SYPRO Orange (Invitrogen), and 10 μM macrodomain protein in buffer containing 20 mM HEPES, NaOH, pH 7.5 and various concentrations of ADP-ribose or hit compounds were mixed on ice in 384-well PCR plate (Roche). Fluorescent signals were measured from 25 to 95 °C in 0.2 °C/30/Sec steps (excitation, 470-505 nm; detection, 540-700 nm). The main measurements were carried out in triplicate. Data evaluation and T_m_ determination involved use of the Roche LightCycler® 480 Protein Melting Analysis software, and data fitting calculations involved the use of single site binding curve analysis on GraphPad Prism. The thermal shift (ΔT_m_) was calculated by subtracting the T_m_ values of the DMSO from the T_m_ values of compounds.

### AlphaScreen (AS) Assay

The AlphaScreen reactions were carried out in 384-well plates (Alphaplate, PerkinElmer, Waltham, MA) in a total volume of 40 μL in buffer containing 25 mM HEPES (pH 7.4), 100 mM NaCl, 0.5 mM TCEP, 0.1% BSA, and 0.05% CHAPS. All reagents were prepared as 4X stocks and 10 μL volume of each reagent was added to a final volume of 40 μL. All compounds were transferred acoustically using ECHO 555 (Beckman Inc) and preincubated after mixing with purified His-tagged macrodomain protein (250 nM) for 30 min at RT, followed by addition of a 10 amino acid biotinylated and ADP-ribosylated peptide [ARTK(Bio)QTARK(Aoa-RADP)S] (Cambridge peptides) (625 nM). After 1h incubation at RT, streptavidin-coated donor beads (7.5 μg/mL) and nickel chelate acceptor beads (7.5 μg/mL); (PerkinElmer AlphaScreen Histidine Detection Kit) were added under low light conditions, and plates were shaken at 400 rpm for 60 min at RT protected from light. Plates were kept covered and protected from light at all steps and read on BioTek plate reader using an AlphaScreen 680 excitation/570 emission filter set. For counter screening of the compounds, 25 nM biotinylated and hexahistidine-tagged linker peptide (Bn-His_6_) (PerkinElmer) was added to the compounds, followed by addition of beads as described above.

### Fluorescence Polarization (FP) Assay

The FP assay was performed in buffer containing 25 mM Tris pH7.5, NaCl 50 mM, 0.025% TritonX-100. All reagents were prepared as 2X stocks and 10 μL volume of each reagent was added to a final volume of 20 μL. Compounds were preincubated with His-Macrodomain proteins (4 μM) for 30’, RT in black 384 well plates (Corning 3575 plates), followed by addition of 50 nM of fluorescein labeled ADP-ribosylated peptide [5Flu-ARTKQTARK(Aoa-RADP)S]. After mixing for a minute, the plate was incubated at 25°C, protected from light and fluorescence polarization was read after 30 minutes, 1h and 2h using a plate reader.

### Gel-based Inhibition of Mono-ADP-ribosylhydrolase activity (de-MARylaion)

PARP10-CD protein was auto-MARylated through incubation for 20 minutes at 37°C with 1 mM final concentration of β-Nicotinamide Adenine Dinucleotide (β NAD^+^) (Millipore-Sigma) in a reaction buffer (50 mM HEPES, 150 mM NaCl, 0.2 mM DTT, and 0.02% NP-40). MARylated PARP10 was aliquoted and stored at -80°C. To test the ability of identified compounds for their ability to inhibit MARylation activity of Mac1, we first incubated each compound with purified SARS-CoV-2 Mac1 in the reaction buffer (50 mM HEPES, 150 mM NaCl, 0.2 mM DTT, and 0.02% NP-40) at 37°C for 30 min. Then, MARylated PARP10-CD was added to this mixture solution and further incubated for 30 min at 37°C. The reaction was stopped with addition of 2X Laemmli sample buffer containing 10% β-mercaptoethanol. Protein samples were heated at 95°C for 5 minutes before loading and separated onto SDS-PAGE cassette (Thermo Fisher Scientific Bolt™ 4-12% Bis-Tris Plus Gels) in MES running buffer. For immunoblotting, the separated proteins were transferred onto polyvinylidene difluoride (PVDF) membrane using iBlot™ 2 Dry Blotting System (ThermoFisher Scientific). The blot was blocked with 5% skim milk in 1xPBS and probed with the anti-mono-ADP-ribose binding reagent/antibody MABE1076 (α-MAR), and anti-GST tag monoclonal antibody MA4-004 (ThermoFisher Scientific). The primary antibodies were detected with secondary anti-rabbit and anti-mouse antibodies (LI-COR Biosciences). All immunoblots were visualized using Odyssey^®^ CLx Imaging System (LI-COR Biosciences). The images were quantitated using the LI-COR Image Studio software.

### ADP-ribosylhydrolase assay

The recently published assay, ADPr-Glo, was used to examine the impact of our top hit compounds on SARS-CoV-2 enzymatic activity (33). Briefly, the compounds were preincubated with SARS-CoV-2 Mac1 (2 nM) and NudF (125 nM) at ambient temperature for 30 min prior to the addition of MARylated PARP-10 derived substrate. The substrate (20 µM) was then incubated with the SARS-CoV-2 Mac1 and NudF at ambient temperature for 30 min. The reaction products were measured with AMP-Glo. Reactions without macrodomains were performed in parallel as a negative control. Luminescence signal was converted to AMP concentration via interpolation from an AMP standard curve. Data plotted are AMP generated by the macrodomain and NudF, subtracted by AMP generated from NudF alone. Inhibition percentages were calculated and non-linear regression analysis was performed in GraphPad Prism.

### A FRET based binding assay and inhibitor profiling

FRET method was utilized for the profiling of MCD-628 a panel of human and viral macrodomains to determine their specificity (30,36). The assay is based on the site-specific introduction of cysteine-linked mono-ADP-ribose to the C-terminal Gαi peptide (GAP) by Pertussis toxin subunit1 (PtxS1) fused to YFP. To generate the FRET signal ADP-ribosyl binders were fused to CFP. Samples were prepared in the assay buffer (for most binders; 10 mM Bis-Tris propane pH 7.0, 3 % (w/v) PEG 20,000, 0.01 % (v/v) Triton X-100 and 0.5 mM TCEP), (for TARG1; 10 mM Bis-Tris propane pH 7.0, 150 mM NaCl, 0.01 % (v/v) Triton X-100 and 0.5 mM TCEP), (for PARG; 10 mM Bis-Tris propane pH 7.0, 25 mM NaCl, 0.01 % (v/v) Triton X-100 and 0.5 mM TCEP) in a 384-well black polypropylene flat-bottom plates (Greiner, Bio-one) with 10 µL reaction volume per well. The reactions consisted of 1 µM CFP-fused binders and 5 µM MARylated YFP-GAP. Reactions were excited at 410 nm (20 nm bandwidth), while the emission signal was measured at 477 nm (10 nm bandwidth) and 527 nm (10 nm bandwidth). Afterwards, blank was deducted from the individual values and the radiometric FRET (rFRET) was calculated by dividing the fluorescence intensities at 527 nm by 477 nm. Compound was dispensed with Echo acoustic liquid dispenser (Labcyte, Sunnyvate, CA). Dispensing of larger volumes of the solutions was carried out by using Microfluidic Liquid Handler (MANTIS®, Formulatrix, Beford, MA, USA). Measurements were taken with Tecan Infinite M1000 pro plate reader.

### Computational modeling

Hit compounds were docked into the ADPr-bound (6WOJ), 3 unique unbound conformations (7KR0, 7KR1, 6WEY) and two small molecule bound (5RSG, 5RTT) structures of SARS-CoV-2 Mac1 (35,39,40). The proteins and ligands were prepared using Schrodinger Maestro and were subsequently docked using Glide with XP precision, analog compounds **6, 7, 8**, and **9** were re-docked using a core constraint to a high scoring, intuitive pose of compound **7**, and high scoring poses were subjected to a Prime MM-GBSA minimization, allowing flexibility for any residue within 5 Å of the ligand (41-46).

## ACKNOWLEDGEMENTS

ARF would like to the the KU Synthetic Chemistry Core facility for help in ordering and confirming several of these compounds. DVF would like to acknowledge McDaniel College Student-Faculty Summer Research Fund, the Jean Richards Fund, the Schofield fund, and the Scott and Natalie Dahne fund. DVF would also like to acknowledge Mr. Kristopher Mason and Vaccitech for the use of their mass spectrometer. LL would like to acknowledge the use of the facilities of the Biocenter Oulu Structural Biology core facility, a member of Biocenter Finland, Instruct-ERIC Centre Finland and FINStruct. This research was funded by National Institutes of Health (NIH) grants P20 GM113117, P30GM110761, a CTSA grant from NCATS awarded to the University of Kansas for Frontiers: University of Kansas Clinical and Translational Science Institute (#UL1TR002366), and University of Kansas start-up funds to ARF, and by Sidrid Jusélius foundation grant to LL, and Johns Hopkins Bloomberg School of Public Health Discretionary Fund to AKLL. The contents are solely the responsibility of the authors and do not necessarily represent the official views of the NIH or NCATS.

---

**Table S1:** Compound docking scores

| **Compound** |  | **glide** |  |
| --- | --- | --- | --- |
| **#** | **docking score** | **emodel** | **Mac1 Structure** |
| 1 | -7.857 | -81.695 | 6WOJ |
| 6 | -4.563 | -50.414 | 7RK0 |
| 7 | -3.338 | -34.249 | 7RK0 |
| 8 | -4.064 | -57.015 | 7RK0 |
| 9 | -4.287 | -61.816 | 7RK0 |
| 10 | -4.806 | -59.941 | 6WEY |
| 11 | -5.306 | -58.757 | 6WEY |

**Figure S1.**
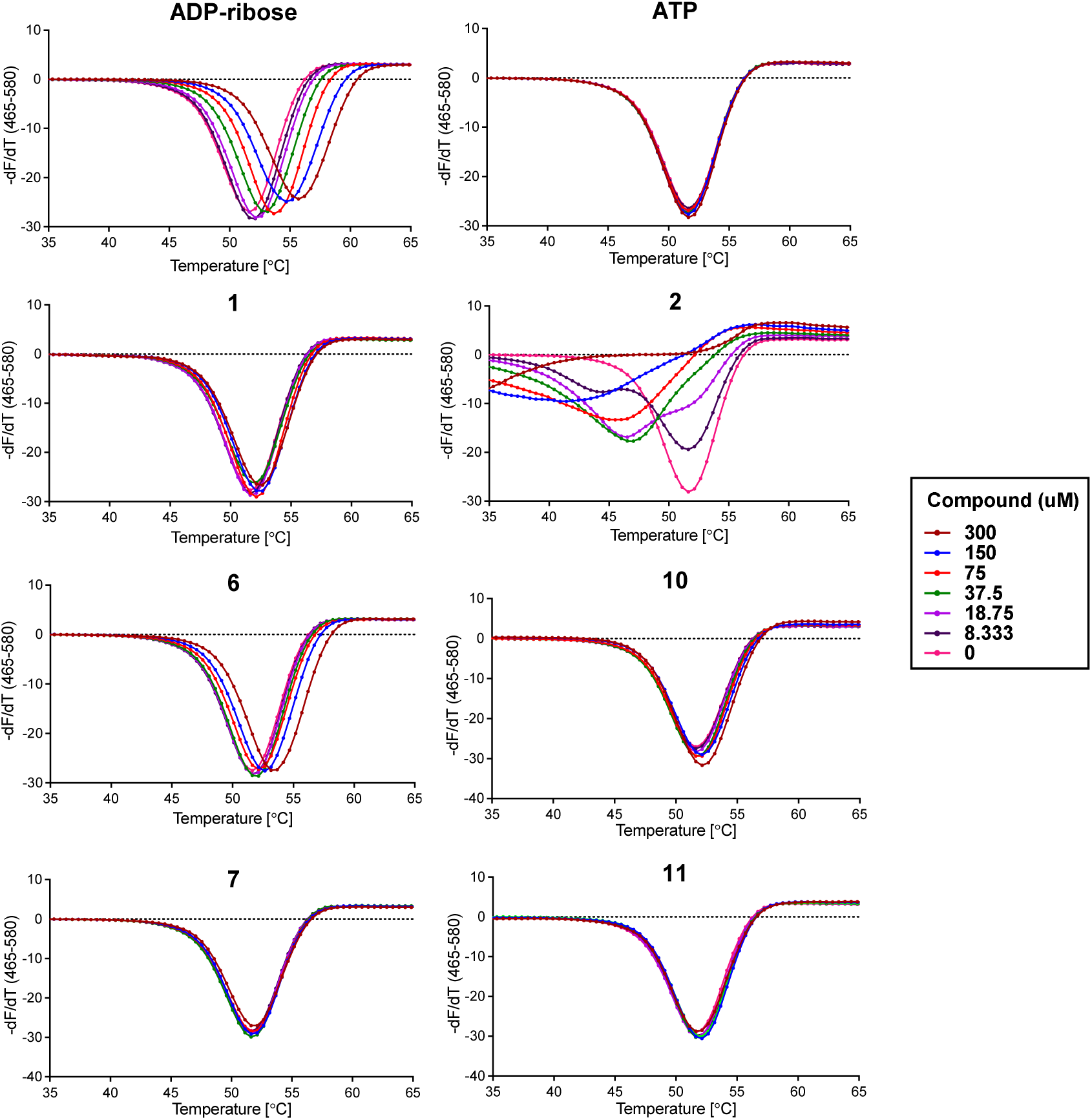
Thermal stability of SARS-CoV-2 Mac1 after incubation with hit compounds. The top 4 hit compounds were tested for their ability to increase the thermal stability of SARS-CoV-2 Mac1 in a differential scanning fluorimetry assay (DSF). Thermal profiles are shown for each compound at different concentrations.

**Figure S2.**
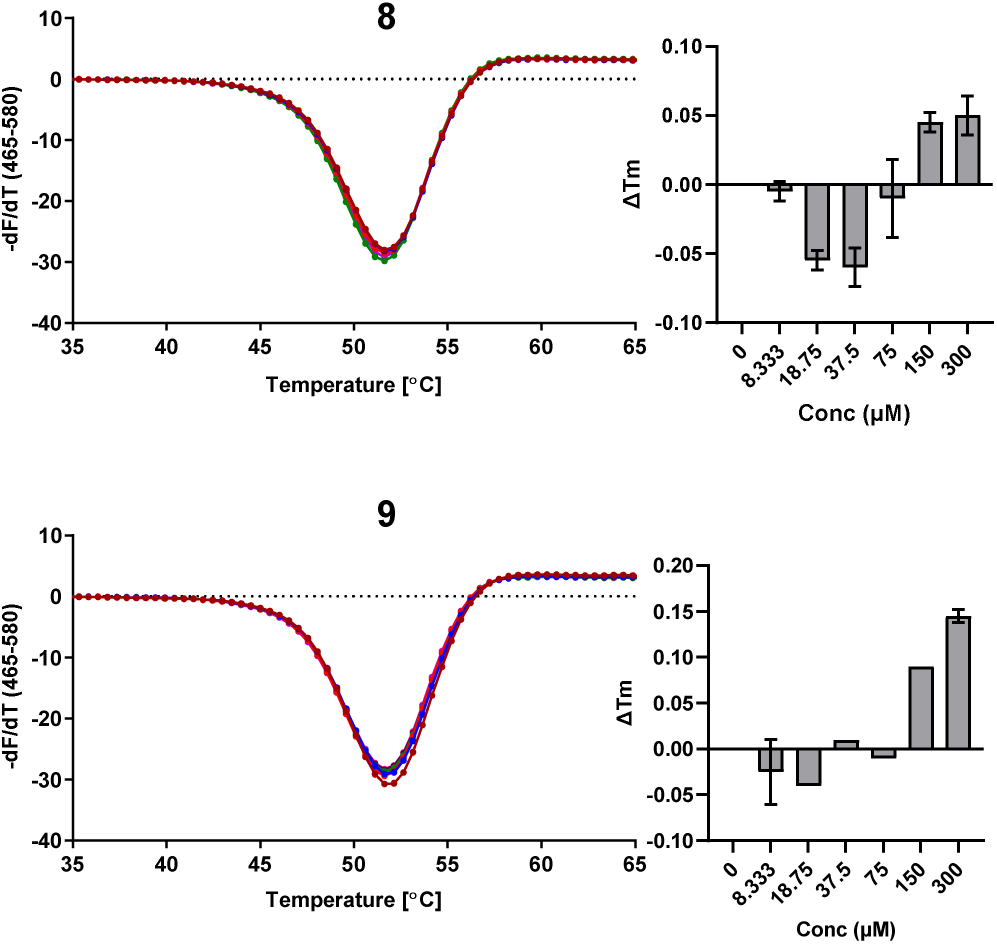
Thermal stability of SARS-CoV-2 Mac1 after incubation with analog compounds. Two analogs of FS2MD-006 are shown here for their ability to increase the thermal stability of SARS-CoV-2 Mac1 in a differential scanning fluorimetry assay (DSF). Thermal profiles and change in T_m_ are plotted for each compound at different concentrations.

